# TSD: A computational tool to study the complex structural variants using PacBio targeted sequencing data

**DOI:** 10.1101/474445

**Authors:** Guofeng Meng, Ying Tan, Yue Fan, Yan Wang, Guang Yang, Gregory Fanning, Yang Qiu

## Abstract

The PacBio sequencing is a powerful approach to study the DNA or RNA sequences in a longer scope. It is especially useful in exploring the complex structural variants generated by random integration or multiple rearrangement of internal or external sequences. However, there is still no tool designed to uncover their structural organization in the host genome. Here, we present a tool, TSD, for complex structural variant discovery using PacBio targeted sequencing data. It allows researchers to identify and visualize the genomic structures of targeted sequences by unlimited splitting, alignment and assembly of long PacBio reads. Application to the sequencing data derived from an HBV integrated human cell line(PLC/PRF/5) indicated that TSD could recover the full profile of HBV integration events, especially for the regions with the complex human-HBV genome integrations and multiple HBV rearrangements. Compared to other long read analysis tools, TSD showed a better performance for detecting complex genomic structural variants. TSD is publicly available at: https://github.com/menggf/tsd

## INTRODUCTION

Structural variations (SV), such as copy number variations, inversions and translocations, are commonly observed in genomes (Feuk *et al*. 2006). In human, SVs exist in approximately 13% of the genome in the normal population (Sudmant *et al*. 2015). Some of these SVs contribute to the phenotype diversity and susceptibility to diseases (Brandler *et al*. 2016; Truty *et al*. 2018), which draws more attention in disease studies. Complex SVs can also be observed in the genome with genetics instability (Weischenfeldt *et al*. 2013; Lupski 2015), virus integration or transgenic modification (Zhao *et al*. 2016a; Meng 2018), which may generate complex rearrangement of external DNA sequences and random integration in the host genome.

The next-generation sequencing (NGS) technologies have been widely used for SV discovery Abel and Duncavage (2013); Tubio (2015). In such studies, NGS platforms typically generate millions of short reads, ranging from 50 to 300 bp long. The SV discovery is performed by the analysis to the short reads deriving from the SV regions, such as discordant paired-end reads, split reads and sequencing depth information (Zhao *et al*. 2016b). However, NGS-based SV discovery is limited by the read length, especially for long complex SVs, which makes the more detailed information of the complex SVs usually blind to computational algorithms. Therefore, most of the NGS approaches primarily focus on low-complexity copy number variants or rearrangements.

The advantage of the third-generation sequencing technologies, such as the PacBio sequencers released by Pacific Bioscience, which generate reads up to 60 kbp long (Rhoads and Au 2015), has been emerging as a powerful approach to study the genome in a longer scope. However, the reads generated from the PacBio sequencer are error-prone, especially the indel errors (Rhoads and Au 2015). It challenges the SV discovery using the tools designed for NGS. Many computational methods have been developed for long read, e.g. *de novo* assembly, isoform studies and other applications (Koren *et al*. 2016; Chin *et al*. 2013; Chaisson and Tesler 2012; English *et al*. 2015). However, most of these tools are not designed or optimized for the genomic regions with complex structure, such as complex chromosome relocation, integration and rearrangements. Meanwhile, the throughput and cost of PacBio platform limit its application only to small-sized genome, e.g. bacteria’s genome Ferrarini *et al*. (2013); Liao *et al*. (2015). Targeted sequencing has been widely applied in studies by capturing only the interest region. When it comes to long reads, how to use such redundant reads for an accurate SV discovery also challenges the data analysts.

Here, we present a tool, TSD, for identifying and visualizing the structural variants using PacBio targeted sequencing data. It is specially designed for the DNA regions with complex integration and rearrangement by allowing multiple rounds of splitting and mapping of the long PacBio reads. The genomic organization structure of targeted sequences is recovered by assembling the mapped PacBio fragments. TSD is applied to a PLC/PRF/5 cell line, which contains complex HBV rearrangement and integration events, and identified 9 HBV integration events. Evaluation suggests that TSD has an equal or better performance in discovering the structure of SVs than existing tools, especially when the targeted sequences have complex structure in the genome.

## MATERIALS AND METHODS

### Targeted sequencing on PacBio Sequel of PLC/PRF/5 cell line

The targeted regions with HBV integrations in PLC/PRF/5 cell genome were sequenced on PacBio Sequel SMRT system. Briefly, genomic DNA was extracted from PLC/PRF/5 cell (ATCC, CRL-8024) using PureLink Genomic DNA Kit (Invitrogen, CAT K182002) followed by random fragmentation to 5-9 kbp long. Fragments containing HBV sequence were captured and enriched using Roche NimbleGen’s SeqCap EZ enrichment technology with customized HBV specific probes. SMRTbell library was prepared according to the manufacturer’s guidelines and sequenced on PacBio Sequel by GENEWIZ Company. The quality control and processing of raw PacBio reads were performed using SMRT Link v5.1.0. The subreads are used as the initial input of TSD analysis.

### Simulated PacBio reads

We generated a set of simulated PacBio reads by randomly extracting the genomic DNA fragments and connecting them to form complex SVs. In this process, the genomic fragments were set to have a random length ranging from 500 bp to 2000 bp. We checked the genomic annotation for the selected fragments and found that they covered diverse regions in human genome, including the coding regions and low-complexity repeat regions. Finally, 5,000,000 genomic fragments were collected and connected in a random way to form 1,000,000 PacBio reads. Each simulated PacBio read carried 3-7 fragments. To simulate the error-prone feature of PacBio reads, indel errors were also added to the reads in a random way by keeping the error ratio to be about 15%. The simulated PacBio reads were used to evaluate the accuracy of alignment mapping and assembly of long PacBio reads.

### Reference index for targeted sequences

The PacBio reads are mapped to reference genome using the Burrows-Wheeler Aligner (`BWA-MEM`) (Li and Durbin 2009). For the data with external sequence, e.g. virus or transgenic vector, supports two ways to build the reference index. One way is to treat the external sequences as the extra chromosomes and build the reference index as a whole. The second way is to build the reference index for both genome sequences and targeted sequences, respectively. In most cases, these two ways have no difference to the analysis results. However, when the targeted sequences are derived from the genome or are homologous to the host genome sequence, e.g. the LINE repeats in the human genome, the corresponding genome regions should be masked using the tools like `RepeatMasker` (Tarailo-Graovac and Chen 2009) or manually replacing the the corresponding nucleotides as “N”s. Otherwise, it will confuse `TSD` in the final output.

### Long reads alignment to the genome

The long PacBio reads are firstly aligned to reference sequences using “bwa mem −x pacbio” command. If the long reads are partially mapped and unmapped part is longer than a minimum length (e.g. 200 bp), the unmapped sequences are further cut out for next-round alignment. This process is repeated for multiple times until no new mapping can be generated. In this way, the PacBio reads with complex structure are represented as a line of mapped fragments.

### Fragment similarity and clustering

The PacBio read derived from the complex SVs may be comprised of multiple mapped DNA fragments, which usually have different origins and organization directions. We used three values to record each mapped DNA fragments: chromosomes, starting points and ending points. Considering the fact that PacBio sequencing data were error-prone, the mapped fragments were clustered into consensus fragments (CFs). In this process, a similarity-based clustering algorithm was applied to cluster the DNA fragments from the same chromosome with the similar starting and ending points. In this way, the PacBio reads were further transformed into a line of connected CFs.

For two CFs chrX:f1-t1 and chrX:f2-t2, their similarity is measured by two scores, S1 and S2. Assuming two CFs have a overlap of s bp, S1 and S2 are defined as

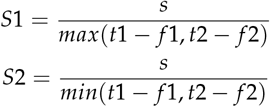

In this work, we used *S*1 > 0.8 as the cutoff to determine if the CFs were derived from the same DNA regions. Alternatively, If the fragments were the first or the last one of PacBio reads, the less strict cutoff *S*2 > 0.8 was used.

### Local alignment of consensus fragments

To determine if two PacBio reads were originated from the same SV, local alignment of PacBio reads was performed by a modified dynamic programming (DP) algorithm. In this algorithm, the CFs instead of nucleotides were used to calculate the matching scores. Two PacBio reads A and B have CFs of {*a*_1_, *a*_2_,..,*a_m_*} and {*b*_1_, *b*_2_,..,*b_n_*}, respectively. A matrix *M*_*m*+1, *n*+1_ was constructed and initialized with 0. The element values of *M*_*m*+1, *n*+1_ were further determined based on the matching status of read A and B. It is described as:

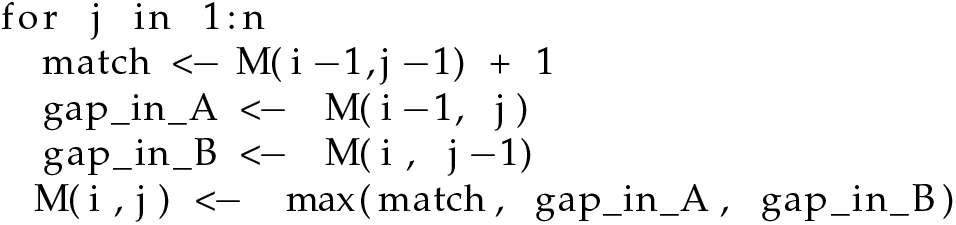

The final alignment of read A and B was tracked back in matrix *M*_*m*+1, *n*+1_. Read A and B would not be treated as reads from the same SVs if any mismatch was observed. Gaps were allowed only when the gaps located in unmapped regions, which were usually resulted from low sequencing quality.

### Seed-based assembly of PacBio reads

Targeted sequencing can generate redundant reads to the same regions. We applied a seed-based clustering method to group the PacBio reads. The long PacBio reads were firstly ranked based on their sequence length or fragment number. The longest read was selected as a seed to assemble the PacBio reads. The other reads were aligned against the seed. If one read was similar with the seed and within the mapping range of the seed, this read would be assigned as a supporting read to the seed. If the read was partially overlapped with the seed, the seed would be extended to construct a new seed. The final seed and its supporting reads were reported as the final outcome.

### Visualization to the structural organization of targeted sequences

The DNA fragment plots are generated using R package to visualize their organization structure of targeted sequences. The plot included multiple information, including (1) the DNA fragments annotated with their origins and the corresponding location; (2) the organization direction of DNA fragments in host genome; (3) the information of supporting reads. Each SV region is reported by a single plot. Users are allowed to modify the output plots by passing the R parameters.

## RESULTS AND DISCUSSION

### Identify the complex structure of targeted sequences

TSD is designed to identify the organization structure of complex SVs, which may be generated by genomic instability, transgene integration, virus infection or even just the low complex genomic regions. As showed in Figure 1(a), such regions may be comprised of multiple fragmented DNA pieces with different origins. To find the detailed information of inserted DNA sequences, it needs computational efforts to mapping and assembling the DNA fragments.

**Figure 1.**
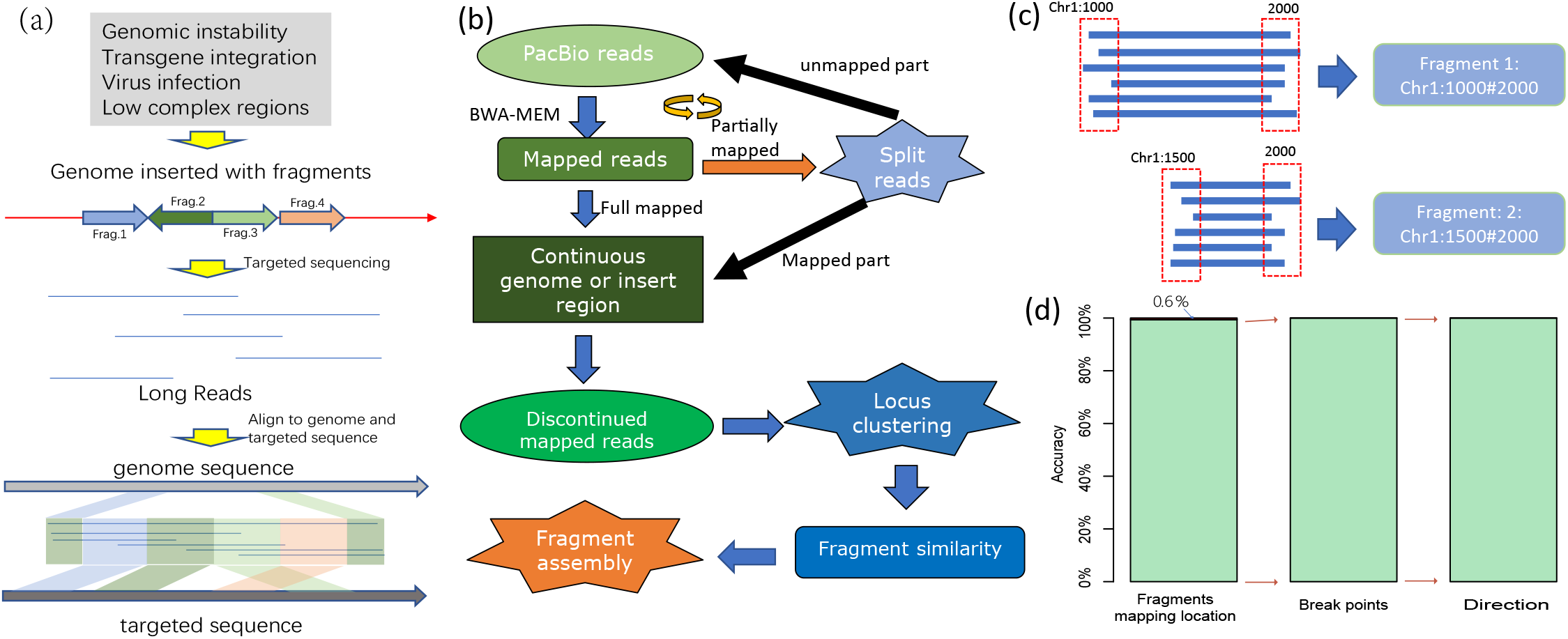
The flowchart of TSD and its evaluation. (a) TSD is designed to identify the structural organization of complex SVs. In this exemplary demonstration, four pieces of external DNA sequences (DNA fragments from the targeted sequences) rearrange and integrate into the host genome. TSD is used to identify their origins, rearrangement and integration location in the host genome. (b) The flowchart of TSD in PacBio reads mapping and assembly, where the long reads are split into mappable fragments and then assembled into readable SVs. (c) The reads mapped to the same locations are clustered together to build consensus fragments. (d) Evaluation using stimulated PacBio reads. 99.4% of reads are correctly mapped to human genome and 100% of stimulated SVs are recovered accurately for both break point location and direction.

Figure 1(b) summarizes the flowchart of TSD in identifying the complex SV structure using PacBio sequencing data. The main idea is that the long PacBio reads are split into mappable fragments and the SV structure are recovered by computational assembly of the mapped DNA fragments (see Figure 1(c)). Compared with the existing tools, TSD has several features: (1) TSD is able to identify the SV structure of any complexity. (2) TSD is designed to analyse targeted sequencing data, where interest sequences are captured or enriched with sequence-specific probes, and TSD can utilize the redundant reads for accurate SV discovery. (3) TSD can display the full structural profile of the complex SVs in the form of plots.

To evaluate its performance, we generated the simulated PacBio reads, including 1,000,000 reads consisting of 5,000,000 genomic fragments. These reads covered diverse genomic regions, including tandem repeats, interspersed repeats and coding regions. In the evaluation to the mapping ability, TSD accurately identified the genomic location of 99.4% DNA fragments (see Figure 1(d)). By checking the fragments with wrong mapping location, we found that all of them were from the repeat regions, especially satellite DNA. These errors were mainly caused by the error-tolerating parameter setting of `BWA` to PacBio reads in alignment, which made `BWA` less specificity to the large tandem repeating DNA. In the evaluation to the assembling ability, TSD recovered the SV structure at an accuracy of 100% when only using the reads with the accurate mapping location. In our evaluation, the genomic structure included both breakpoint position and direction. Both results indicate that TSD can accurately identify the organization structure of complex SVs.

### TSD discovers HBV integration events in PLC/PRF/5 cells

TSD was applied to study HBV integration events in HBV infected PLC/PRF/5 cells. After DNA fragmentation, the HBV-specific probes were used to capture and enrich the DNA pieces that carry HBV sequences. Using 2 million subreads as input, TSD discovered 9 HBV integration events, including the HBV rearrangement, HBV integration and genomic relocation. In total, 12 chromosomes got involved. Figure 2(a) illustrated an exemplary HBV integrated region. This region was extremely complex, consisting of 6 fragments, including two HBV rearrangement generated by linking HBV:2656 to HBV:1446 and by linking HBV:2876 to HBV:2694. The left side of HBV sequence was integrated into chr1:143240209 and the right side was integrated in chr8:35446393. We noticed that such a HBV rearranged sequence was about 2400 bp and no single PacBio read covered the whole region. It was recovered by assembling the PacBio reads. As targeted sequencing was performed, redundant reads were mapped to this region, including 538 reads mapped with consistent location and direction with seed read. In this example, each rearrangement and integration site was also supported by multiple reads. However, we also noticed that more reads were enriched in the HBV regions and their abundance was correlated with the length of HBV sequences. Figure 2(b) showed the genomic organization of other HBV integration events. Like the example in Figure 2(a), we observed complex genomic organization of HBV sequences in human genome. There were only three integration events without HBV rearrangement.

**Figure 2.**
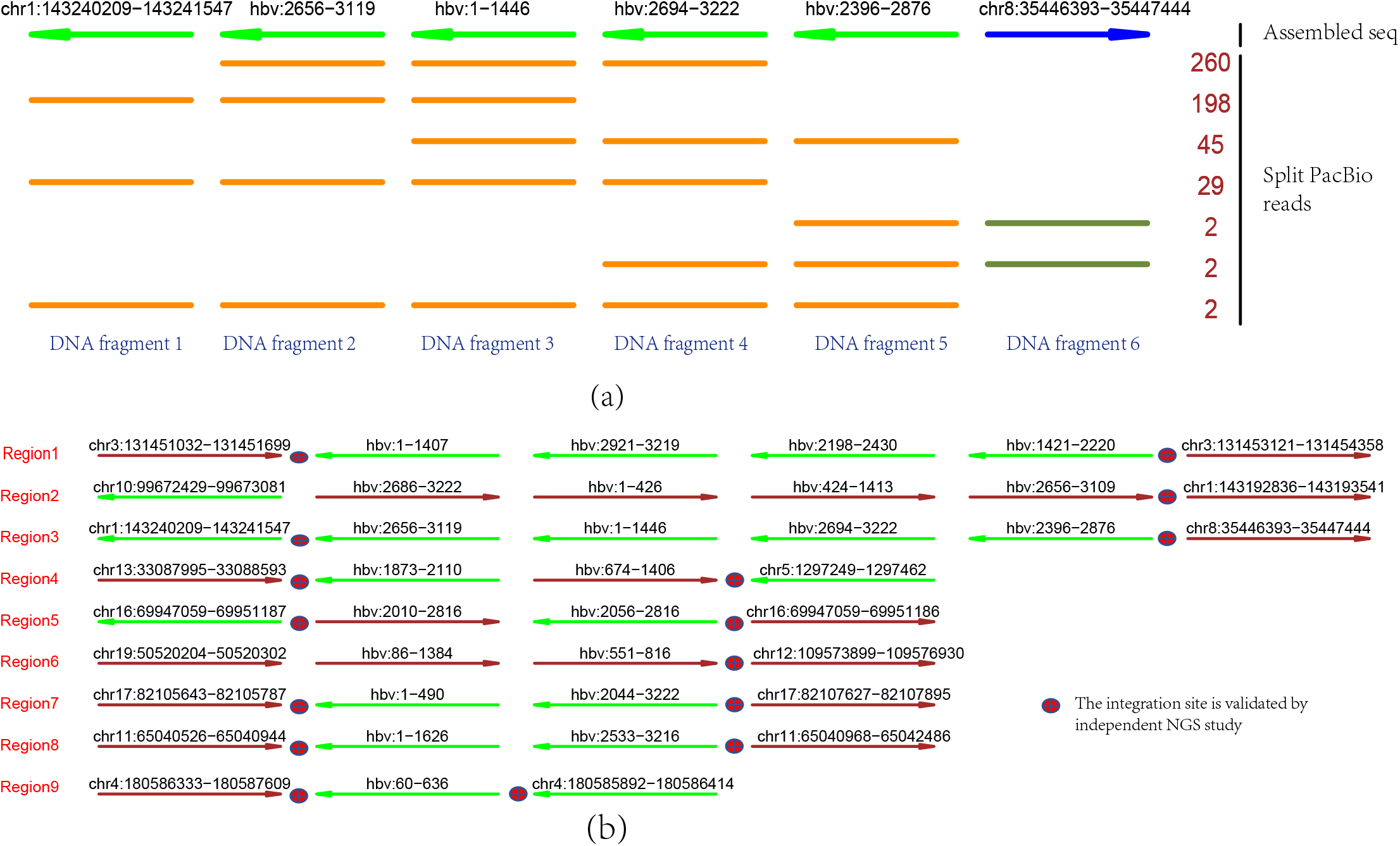
TSD discovers the HBV integration events in the PLC/PRF/5 cells. (a) A genomic region identified by TSD with HBV integration event, which consists of 6 DNA fragments. The first line indicates the assembled HBV integrated region and the other lines indicate the PacBio reads derived from this region. Each row can represent multiple PacBio reads if they have the same fragments composition; the number (right side) counts the PacBio reads. The line colors and arrows indicate the mapped strands: darkgreen: forward; brown: reverse stand. (b) The HBV integration events discovered by TSD using PacBio sequencing data. TSD identified 9 HBV integrated regions, including multiple regions with complex HBV rearrangements. Most of the integration sites are validated by NGS study.

To validate the TSD analysis results, we performed next generation sequencing (NGS) on the same cell line. The integration and rearrangement sites were discovered by analyzing the reads crossing the break points. In Figure 2(b), we marked the integration events validated by NGS analysis results. We found that, all the identified break points could also be discovered by NGS data with the same location and direction. Overall, our results indicated a good confidence for TSD analysis results.

### Evaluation with other tools

We evaluated the performance of TSD by comparing it with the analysis results of HGAP, a *de novo* assembly tool developed by Pacific Biosciences company (Chin *et al*. 2013). Using the same PacBio sequencing data from HBV infected PLC/PRF/5 cells as the input, HGAP outputs the assembled DNA sequences. As the PacBio reads were generated after probe-specific enrichment to the HBV sequence, the output of HGAP were long DNA fragments, including 59 DNA sequences. We mapped them to human genome and HBV sequences with `blast`, respectively. We found one complete integration and several partial integration events. The complete integration event was featured by mapping to both human genome and HBV sequence, allowing inferring its integration location and direction. It started from chr17:82105786 and ended at chr17:82107610 (see Region 7 in Figure 2(b)). There were also partial integration events on chromosome 3 (Region 1, left), 4 (Region 9, left), 5 (Region 4, right) and 11 (Region 8 left), where only either the left integration site or the right integration site was discovered. By checking the integration location for both complete and partial integration events, we found that TSD discovers all these events at the exactly same genomic and HBV locations. Advantageously, TSD also discovered the corresponding matched site for all the partial integration events. Overall, TSD is better at discovering the complex genomic organization structure of targeted sequence than the *de novo* assembly tool.

Another evaluation is performed with Sniffles, a tool designed for SV discovery (Sedlazeck *et al*. 2018). Using the PacBio sequencing data, Sniffles identified many SVs. After filtering the SVs without HBV integration or rearrangement and the ones with low reads coverage, we identified 16 HBV integration sites. Among them, 14 out of them were also discovered by TSD prediction. By checking the NGS analysis results, all of 14 overlapped integration sites could be validated as true positive discovery while the remnant two integration sites didn’t. Compared to the output of TSD, all the prediction reported by Sniffles have been identified by TSD, which is consistent to the algorithm setting of TSD. Meanwhile, the emphasis of TSD is to generate the full profile of complex genomic structure. On the contrary, extra analysis is needed to generate a similar profile from the output of Sniffles.

In Table 1, we summarize the evaluation results of three tools and NGS analysis. By checking the HBV integration events, we observed several reasons that accounted for the output difference. For example, the long SVs, especially the ones without single long read covering the whole regions, need the computational tools to split and assemble the reads for a full profile of SVs. This will be affected by the sequencing depth and quality, computational analysis accuracy and existence of reference genome. For the long complex SVs, TSD can achieve a better performance. Meanwhile, the geonomic location of SVs can also affect the SV discovery. The repeat regions can confuse the alignment tools for wrong genomic locations.

**Table 1.** Evaluation with HGAP and Sniffle.

|  | HGAP |  | Sniffles |  | TSD |  | NGS |  |
| --- | --- | --- | --- | --- | --- | --- | --- | --- |
|  | Int. | Rea. | Int. | Rea. | Int. | Rea. | Int. | Rea. |
| Region1 | left | no | right | no | both | yes | right | - |
| Region2 | no | no | both | no | both | yes | both | - |
| Region3 | no | no | right | no | both | yes | both | - |
| Region4 | right | no | both | yes | both | yes | both | - |
| Region5 | no | no | both | yes | both | yes | right | - |
| Region6 | no | no | both | yes | both | yes | both | - |
| Region7 | both | yes | both | yes | both | yes | both | - |
| Region8 | left | yes | both | yes | both | yes | both | - |
| Region9 | left | yes | both | yes | both | yes | both | - |

Int.: Integration sites; Rea: Rearrangement sites

Overall, our evaluation suggests that TSD can achieve an equal or better performance in identifying the organization structure of complex SVs, especially when the SVs consist of multiple rearrangement events.

### Command-line implementation of TSD

TSD is coded in Perl language and can be implemented in a command-line way. Before usage, TSD requires some preliminary works. Firstly, `BWA` must have been installed and its location added into the $PATH variable in Linux system. Secondly, the genome index has been built using the “bwa index” command.

To simplify the usage, TSD only has two mandatory inputs: “-s” and “-G”, where the former is path location of the PacBio reads file in the format of “*fastq” or “*.fq” and “-G” specifies the prefix of BWA genome index. Users can use “-i” to specify the targeted sequence. If the targeted sequence is foreign DNA, e.g. virus sequence, the “-i” option is mandatory for the reason that TSD cannot assemble the targeted sequences in a *de novo* way. The whole analysis can be done in a way like:

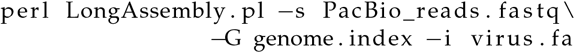

The other parameters are optional. By default, TSD is optimized for the targeted sequencing data, which allow each SV supported by multiple PacBio reads. When the input data is not generated by targeted sequencing, “-r” and “-o” option should be modified, e.g. “-r 1” and “-o 1”, to capture the regions with low sequencing depth. However, this may generate a redundant output. To organize the output, users can use “-d” option to set the directory to store the output and temporary files, which also allows continuous analysis (see software manual for detailed information).

